# Coupling Terrestrial Laser Scanning with 3D Fuel Biomass Sampling for Advancing Wildland Fuels Characterization

**DOI:** 10.1101/771469

**Authors:** Eric Rowell, E. Louise Loudermilk, Christie Hawley, Scott Pokswinski, Carl Seielstad, Lloyd Queen, Joseph J. O’Brien, Andrew T. Hudak, Scott Goodrick, J. Kevin Hiers

**Affiliations:** Tall Timbers Research Station and Conservancy, Tallahassee, FL, USA; USDA Forest Service, Southern Research Station, Center for Forest Disturbance Science, Athens, GA, USA; University of Montana, National Center for Landscape Fire Analysis, Missoula, MT; USDA Forest Service, Rocky Mountain Research Station, Forestry Sciences Laboratory, Moscow, ID, USA

## Abstract

The spatial pattern of surface fuelbeds in fire-dependent ecosystems are rarely captured using long-standing fuel sampling methods. New techniques, both field sampling and remote sensing, that capture vegetation fuel type, biomass, and volume at super fine-scales (cm to dm) in three-dimensions (3D) are critical to advancing forest fuel and wildland fire science. This is particularly true for computational fluid dynamics fire behavior models that operate in 3D and have implications for wildland fire operations and fire effects research. This study describes the coupling of new 3D field sampling data with terrestrial laser scanning (TLS) data to infer fine-scale fuel mass in 3D. We found that there are strong relationships between fine-scale mass and TLS occupied volume, porosity, and surface area, which were used to develop fine-scale prediction equations using TLS across vegetative fuel types, namely grasses and shrubs. The application of this novel 3D sampling technique to high resolution TLS data in this study represents a major advancement in understanding fire-vegetation feedbacks in highly managed fire-dependent ecosystems.

## Introduction

Capturing the processes that influence fire propagation across fuelbeds is the crucial linkage between wildland fire behavior and post-fire effects (Dupuy et al. 2011; Hiers et al. 2009; Linn et al. 2005; Morvan and Dupuy 2001). Yet, characterization of both energy release and fire effects at multiple and appropriate scales remains elusive in providing mechanistic linkages (O’Brien et al. 2018). Capturing multiscale variation in vegetation and representing this complexity as fuel that drives coupled fire-atmospheric interactions remains an unknown challenge.

In general, we know that the spatial patterns of vegetation impacts fire behavior, directly and indirectly. At finer scales (<1m^2^) fuel arrangement, biomass, and type affect fire spread patterns and heat release rate (Hiers et al. 2009; Loudermilk et al. 2012) and at coarser scales (>1m^2^) surrounding vegetation structure and continuity influences air resistance, turbulence (Cochrane 2003; Pimont et al. 2016), and wind field flow throughout the surface fuels and across a stand (Parsons et al. 2017).

Fuels are traditionally represented as two-dimensional abstractions of mass, bulk density, particle density, and surface area that require assumptions that oversimplify fuel elements particularly at scales <1 ha (Hiers et al. 2009). Traditional surface fuel measurements were developed to support coarse grain and point-based fire behavior modeling that did not include a full range of variability or spatial non-uniformity typical of vegetative communities (Hardy et al. 2008). These approaches were designed to report unit averages that give broad landscape-scale assessments of load (Ottmar et al. 2016a; Ottmar et al. 2003). Measuring surface fuels in particular is inherently difficult because they are small--often less than 1-m tall--with litter found below 10 cm (Ottmar et al. 2003). Moreover, shrubs and trees have complex architecture that can influence flow and fire behavior (Parsons et al. 2017).

Traditional measurements of surface fuelbeds include both direct and indirect efforts to supply stand level inputs to large scale fire behavior prediction tools. Common direct measurements are tallies of down woody fuels along planar transects (Brown 1974) coupled with destructive biomass sampling, or “clip plots” (Brown 1981). Indirect methods include visual cover estimates in plots or comparisons with photographs of known fuel loads or types (Keane and Dickinson 2007; Ottmar et al. 2003). These methods provide estimates of gross characteristics— such as fuel load, bulk density, and packing ratios— that are used for predicting fire behavior at the stand or landscape level (Burgan and Rothermel 1984, Reinhardt and Keane 1998, Andrews et al. 2004). They inherently require unrealistic assumptions regarding specific bulk densities for grasses and shrubs (Van Wagner 1968), and are of limited utility for estimating fine-scale fuel heterogeneity that is important for simulating within-stand fire behavior (Linn et al. 2013), and fire effects (Hiers et al. 2009; Loudermilk et al. 2018; Loudermilk et al. 2012; O’Brien et al. 2018).

Current operational vegetation data are represented as fuel models: a set of fuel parameters that characterize a broad spectrum of surface fuelbed properties that are less than 1.83m in height (Rothermel 1972; Scott and Burgan 2005). These fuel models drive the current cadre of operational fire behavior models (e.g. FARSITE; Finney 1998, FLAMAP; Finney 2006; BEHAVEPlus: Andrews et al. 2005). Varner and Keyes (2009) discuss the inherent errors associated with using these fuel models for assessing fuel treatment effectiveness. Noonan-Wright et al. (2013) compared customized fuel models with the existing standard fire behavior fuel models (Scott and Burgan 2005) and found no clear advantage for fuel model customization due to uncertainty of using uncalibrated custom models. Another challenge with the fuel model approach is that fuels are represented as uniform, when in reality they are more often patchy, especially where fuel manipulation through treatments has been performed (Varner and Keyes 2009). To fully understand and model fire behavior (fluid flow, turbulence, flaming front interactions) as a function of fire-atmospheric interactions that function three-dimensionally requires detailed and accurate accounting of fuels also in three-dimensions (Dupuy et al. 2011; Mell et al. 2013; Pimont et al. 2016). Three-dimensional representation of forests characteristics is particularly relevant to surface fire regimes (Hiers et al. 2009; Loudermilk et al. 2012), where fine-scale variation in fuel characteristics have been shown as the important link between structure and function in the U.S. (Glitzenstein et al. 1995; Mitchell et al. 2009; Rebertus et al. 1989; Williamson and Black 1981) and in similar ecosystems globally (Dalgleish et al. 2015; Moreno and Oechel 2012; O’Brien et al. 2008). In these systems, fire behavior and effects are spatially correlated with vegetation structure and patterns at relatively fine scales (<0.25 m^2^; (Hiers et al. 2009; Loudermilk et al. 2012; O’Brien et al. 2016). Previous studies have suggested that this relationship is due to variations in ignition and combustion characteristics of different vegetation types and their influence on the ambient and fire induced wind flows (Fernandes et al. 2004; Hoffman et al. 2016; Linn and Cunningham 2005; Linn et al. 2013). These fire-atmosphere interactions have traditionally been assessed in isolation or at the wrong scales required to advance fundamental understanding of how various factors control fine scale variability in fire behavior and effects. Most managers and researchers use the aforementioned modeling tools (e.g., BEHAVE, FFE-FVS, FCCS) where forest structure and fuels are overgeneralized (Andrews et al. 2005) and ignore the spatial complexity of the fuels and fire-atmospheric feedbacks.

Computational fluid dynamic (CFD) fire behavior models, such as FIRETEC (Linn et al. 2005) and the Wildland Fire Dynamics Simulator (Mell et al. 2009), operate on 3D representations of vegetation and provide new opportunities to understand the underlying mechanisms and interactions driving complex fire behavior. For example, recent studies have utilized these models to gain new insight into the dominant controls of fire spread (Linn and Cunningham 2005; Mell et al. 2009), energy release (Linn et al. 2002; Linn et al. 2005), 3D canopy-mediated flow (Dupuy et al. 2011), and spatial patterns of fuel consumption (Parsons et al. 2011). These detailed physics-based models perform optimally when the three-dimensional nature of the entire fuels complex is represented within the model. It is now possible to characterize finer scale aggregation of interacting vegetation types as discrete 3D “wildland fuel cells,” (*sensu* Hiers et al. 2009) which can be mapped within the surface fuelbed of CFD models.

In the past decade, there have been numerous advances characterizing direct measurements of fuel mass using terrestrial laser scanning (TLS) (Newnham et al. 2015). TLS data excels at measuring fine-scale fuel structure at local scales (<0.25 m^2^ grainsize) for primarily producing estimates of fuel mass and bulk density. These instruments have the ability to collect 3D structural information on objects with less than 1 cm accuracy and precision, which provide a data richness that out-performs typical field methods. Their measurements are used to produce precise fine-scale volumetric estimates that correlate well with biomass and leaf area (Greaves et al. 2015; Loudermilk et al. 2009; Olsoy et al. 2014; Rowell et al. 2016b), are linked to fine-scale fire behavior (Loudermilk et al. 2012), and are used to create fuel height models for fine-fuel types, such as grasses, shrubs, and litter (Rowell and Seielstad 2012; Rowell et al. 2016b). There are examples of the integration of TLS data with Airborne Laser Scanning, structure from motion, and as supplemental training data for improved landscape estimates of fuels (Cooper et al. 2017; Greaves et al. 2017; Rowell et al. 2016a). These approaches now allow for direct, spatially explicit, and quantifiable estimates of local and landscape scale fuel estimates that can be used for fuels planning and eventually operational physics-based fire behavior modeling.

A key limitation in TLS-based fuels characterization to date has been the use of traditional two-dimensional field data for validation; this approach precludes the ability to compare directly from remotely sensed 3-D data. Measuring biomass and structure of surface fuels is complex because neither field measurements nor remote sensing instrumentation have been able to estimate both the 3D distribution and mass of *intermixed* fuel types. And although coupling field data (fuel types, biomass) with remotely sensed data (TLS: volume) would seem to provide the information needed, the disconnect lies in the need for the 3D mass and volume of each fuel type within a fuel patch at the appropriate scale (Hiers et al. 2009). Two-dimensional biomass clip plots (1m^2^) do not link well to structural estimates captured at finer 3D scales (1 cm^3^). Even with spectral information, the data are limited to fuel types found in the sensor’s field of view (Loudermilk et al. 2014), where obstruction biases towards the taller vegetation. Coupling these datasets to estimate, for example bulk density (mass per unit volume), limits the scale of output information to the size of the clip plots, and limits the quality of the outputs by the complexity of the fuelbed matrix (Hiers et al. 2009).

The objective of this study was to integrate a novel fine-scale 3D field sampling technique with TLS-based measurements to better characterize surface fuels, with the assumption that 3D field data should better link with 3D laser data. First, we assessed the variability in fuel mass as a function of fuel type and height class, understanding that currently, there are no fuel types associated with the laser data but we can infer distributions of fuel mass if other structural fuel characteristics are known. Secondly, we provide an analysis of TLS-derived fuel metrics to assess the relationships between fine-scale mass and structural metrics including fuelbed occupied volume, porosity, and surface area.

## Methods

### Study Area

Field measurements and TLS data were collected at the 1,222-ha Pebble Hill Plantation (PHP) in the Red Hills region of southern Georgia, USA (30° 35’N, 84° 20’W, elevation 60-85 m above sea level) as part of a series of prescribed fires conducted through the Prescribed Fire Science Consortium. The Red Hills region has a temperate sub-tropical climate of warm to hot, humid summers and short, mild winters with mean monthly temperatures ranging from 26.8 °C in July to 10.4 °C in January (Arguez et al. 2010). Mean annual precipitation (recorded 21 km to the south at Tall Timbers Research Station, 1878-2010) is 1,359 mm. PHP was utilized as a cotton plantation until 1896 when the property then proceeded to be managed for hunting of the northern bobwhite quail (*Colinus virginianus* L.) which included allowing most agricultural fields to succeed to old-field pine-grasslands and maintaining fire intervals that are typically every one to two years (Robertson and Ostertag 2007). The old-field pine-grassland communities are dominated by shortleaf pine (*Pinus echinata* Mill.) and loblolly pine (*Pinus taeda* L.), where the understory is a continuous layer of grasses, forbs, and hardwoods maintained in a shrub state through frequent fire (Ostertag and Robertson 2007). In sites where no previous farming or tilling occurred, approximately one-third of PHP pinelands, native longleaf pine (*Pinus palustris* Mill.) with a wiregrass (*Aritistida stricta* Michx.) understory is found and is classified as Clayhill Longleaf Woodlands (Carr et al. 2010; Ostertag and Robertson 2007).

In March and April of 2017, a total of 20 clip plots (aka “plots”) were distributed among four burn units at PHP. The plot locations were strategically located to represent the fuel types characteristic of a longleaf pine woodland that is typically burn every 1–3 years. The northwest corner of each plot was monumented with a 1.5 m tall metal pole wrapped with highly reflective retro-tape. After TLS was conducted and before prescribed burning, the plots were sampled and harvested for 3D biomass measurements using the most recent 3D fuels sampling protocol (Hawley et al 2018). This approach uses a voxel sampling framework, which employs an adjustable 3D rectangular sampling frame that allows fuels data to be collected in the field at three different scales—entire plot (0.25 m3), stratum (0.025 m3), down to individual voxels (0.001 m3). The 3D sampling frame outlines the sampling area that is 0.5 m in width by 0.5 m in length by 1 m in height. The frame is subdivided into ten 10 cm vertical sampling strata and each 10 cm stratum contains twenty-five 10 cm3 cells, totaling 250 voxels that are distributed over the frame’s sampling volume of 250,000 cm3 or 0.25 m3.

Within each plot, starting at the highest stratum that contained vegetation, each voxel was sampled for presence/absence of fuel type. Specific to a longleaf pine woodland, the fuel types included 1–10 hour fuels, 100–1000 hour fuels, general pine litter (e. g. shortleaf and/or loblolly pine), wiregrass/bunchgrass, other graminoids, shrubs, volatile shrubs, forbs, pine cones, deciduous oak litter, evergreen oak litter, and longleaf pine litter (see Hawley et al. 2018). Each voxel had the potential to encompass multiple fuel types. Heights of fuels were collected as a function of voxel cell position. Once the fuel types were recorded for each voxel within a stratum, all biomass within the stratum was destructively harvested by clipping and bagging the material. The occupied voxel sampling and biomass collection method was repeated every 10 cm down the frame until mineral soil was reached.

The collected biomass was dried at the United States Department of Agriculture Forest Service, Southern Research Station, Forestry Sciences Laboratory, located in Athens, GA at 70 degree Celsius until the weight of the sample no longer changed. For most material, this required 48 hours of drying time. Some of the heavier fuels required 72 to 96 hours of drying time.

### Terrestrial Laser Scanning

Terrestrial laser scanning was conducted using a RIEGL VZ2000 to collect three-dimensional point clouds at ~5mm point spacing at 15m range. The VZ2000 is a near infrared eye safe laser that is capable of scanning objects at up to 1000m. Laser scanner collection points were established on the four corners of the rectangular burn units, set back a minimum 2.5m from the unit edge. A single 360° scan was collected in the center of the burn unit. Scan parameters were set to sample points at 0.023° frequency at a scan rate 550 kHz per scan. Individual scans were geospatially located using the onboard GNSS L1 GPS receiver that automatically places all points into the desired spatial reference (UTM 16N, NAD83). Data were exported to a LAS file format using the proprietary software RiSCAN Pro (RIEGL, Austria). Fine-scale correction between scans was performed using the freeware CloudCompare (http://www.cloudcompare.org), where identifiable features between multiple scans were used to “stitch” individual scans to the best accuracy possible (~1cm RMSE). Final merged datasets were compared to GPS monuments at the corners of burn units and at the NW corner of each sampled plot to produce a final georeferenced data product. In CloudCompare, all scans were merged into a single dataset and exported into an ASCII text format, with scan variables of x,y,z, and reflectance intensity included.

### Voxelization of TLS Data

Distillation of the three-dimensional TLS point data to relevant scale and metrics used for fuel mapping required conversion into three-dimensional voxel space. For ease and replication, we employed the VoxR package (Lecigne et al. 2014) in R, where voxel domains are established and modeled over relatively large areas at low computational cost to the user. Voxel cell domains were defined as 10cm^3^. This domain size is substantially aggregated enough to derive important metrics for assessing fuel characteristics and sufficiently fine-scale to capture large gaps and variability in the fuelbed. From the voxel analysis, we calculated an occupied volume per plot, following Rowell et al. (*In review*). Additionally, we calculated a porosity and surface area metric for each 10cm^3^ voxel cell using a sub-voxel approach. These metrics are further explained in the following sections.

### Surface Fuel Porosity

Fuel porosity derived from TLS data is a metric similar in practice to packing ratio or as porosity as defined by Anderson (1969). Packing ratio is defined as the fuel load of similar density divided by the fuelbed depth. This metric serves as an estimate of compactness that is critical to explaining how fire propagates through a porous medium. This compactness represents the expected movement of air that affects residence time and combustion intensity (Anderson 1969). As the concept of porosity or packing density from the Anderson (1969) perspective already integrates fuel mass in the calculation, we expect that calculation of TLS-derived porosity would be well correlated with measured fuel mass.

To test this hypothesis, we calculate TLS-based porosity using equation 1:

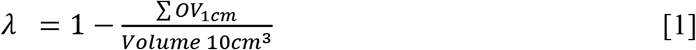

Where, porosity (λ) is the relative proportion of open space resulting from the occupied volume (OV) divided by the total volume of the 10cm^3^ voxel. This assumes that omission and commission errors are the same across voxels. We calculated this definition of fuelbed porosity for the 0-10 cm and 10-20 cm stratum only, as these strata represent where the highest proportion of compact fuels and fuel mass that typically occur in a frequently burned fuelbed (Rowell et al. 2016).

### Voxel-based Surface Area

For fuels sampled above 10 cm, we calculated surface area within each occupied voxel. To estimate surface area of fuel elements at the 10 cm3 voxel domain, the points within each voxel were subset and recalculated using a 3D kernel density estimate via the kde3d function included in the misc3d package in R (Tierney 2015). The kernel density function weights distributions of points to subsequently estimate better isosurfaces that can be used to predict surface area (Feng and Tierney 2008). We used the vcgIsosurface function as part of the Rvcg package in R (Schlager 2017) that represents constant densities of the kernel density function over the limits of the voxel domain. This method used the marching cubes algorithm (Lorensen and Cline 1987), that created a surface through intersecting edges of a volume grid with a volume contour. Where edge intersections occur, a vertex was created. The surface area of the fuel element for the voxel domain was calculated using the vcgArea function within Rvcg, which calculates the surface of the triangular mesh from the isosurface.

### Statistical Analysis

We compared estimates of occupied volume from the voxel sampling method and the TLS-based method with each other and tested the predictive power of each method to estimate total fuel mass for the plots using a leave-one-out-cross-validation (LOOCV) method in the caret package for R (Kuhn 2013). We also analyzed the relationship between measured fuel mass by stratum with maximum porosity for the 0-10 cm and 10-20 cm surface fuel characteristic, and produced a multiple linear model with the max porosity from both strata using LOOCV. We selected the maximum porosity metric as our focus metric, as there was a wider range of estimates of porosity over the mean porosity metric, which is less sensitive to inflections that correspond to variability within the fuelbed (e.g. pine cones, coarse woody debris). Across all strata (1-100 cm heights), we analyzed the relationship between measured fuel mass by stratum and total surface area per stratum and produced a linear model relating fuel mass with total surface area per stratum using the LOOCV method. While investigating the performance of porosity and surface area for predicting fuel mass, it became apparent that porosity was a stronger metric lower in the fuel bed and surface area was stronger higher in the fuelbed. Following systematic testing, we utilized the porosity metric for the 0-10 cm and 10-20 cm height strata and the surface area metric above 10 cm. We then combined the predictions from the maximum porosity and surface area models to produce a new estimate of total fuel mass per plot and produced a linear model between measured and combined fuel mass using LOOCV. We speculated that porosity and surface area perform differentially due to variable compactness of fuels with height. Lower in the fuelbed where most of the litter mass was found, porosity better differentiated between filled and empty space (Rowell et al. 2016).

For all model estimates we assessed accuracy using root mean square error (RMSE) and reported both RMSE and the percent error of the mean measured fuel mass. We also tested the equivalence of the estimates and the measured fuel mass using the equivalence package in R (Robinson and Robinson 2016). The equivalence test was assessed based on a null hypothesis of dissimilarity.

## Results

### Occupied Volume and Mass from TLS Occupied Volume

Total occupied volume was correlated between the measured (observed) data and laser data at the plot scale, explaining 85% of the variability (Adj. R^2^ = 0.85, figure #) with an error of 16% (RMSE = 0.014m^3^). To assess the variability at a stratum level, laser estimated occupied volume was compared with observed occupied volume in each stratum resulting in laser based OV explaining 86% of the variability (Adj. R^2^ = 0.86, **Fig. 1**).Error associated with per stratum analysis was 41% overall (RMSE = 0.003 m^3^) with the mid-values of occupied volume encompassing the greatest range of uncertainty.

**Figure 1.**
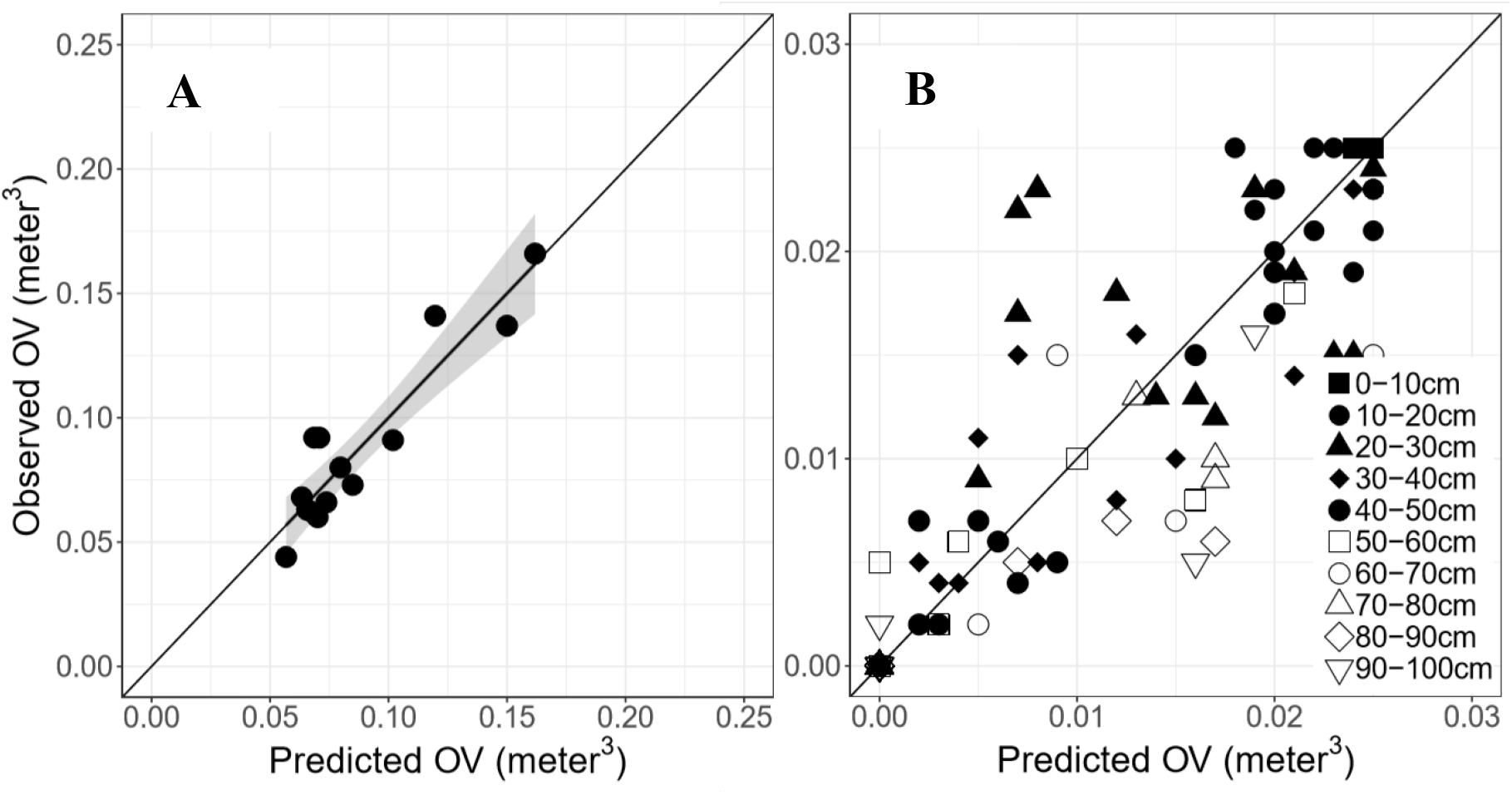
Scatterplots of the TLS-based occupied volume (Predicted OV) with measured occupied volume (observed OV) from the 3D sampling protocol at the plot (A) and stratum (B) level, across all 20 plots. The line illustrates a 1:1 relationship. Note the differences in scale in both axes between A and B.

Measured occupied volume was also correlated to total fuel mass (Adj. R^2^ = 0.27; p-value = 0.033) with a somewhat stronger relationship found between laser estimated occupied volume and measured fuel mass (Adj. R^2^ = 0.44; p-value = 0.005), with associated errors of 33% (RMS E = 43.43 g) and 28.5% (RMSE = 37.69 g), respectively.

### Fuel Mass from TLS Porosity and Surface Area

Fuelbed maximum porosity was linearly correlated with dry-weight biomass in the 0-10 cm and 10-20 cm strata and a multiple linear model explained 89% of the variability (Adj. R^2^ = 0.90, **Fig. 2**). The porosity modeled predictions of mass produced a 20% error (RMSE = 32.0 g) and bootstrap tests of equivalence rejected the null hypothesis of dissimilarity (*P* = 0.025). For fuel features between 10-100 cm, total surface area for each stratum was correlated with dry-weight biomass for wiregrass plus litter fuels (Adj. R^2^ = 0.68, **Fig. 3A**) and shrub fuels (Adj. R^2^ = 0.52, **Fig. 3B**). The model for grass plus forbs fuels yielded an error of 90% (RMSE=3.58 g) and an error of 54% (RMSE = 8.28g) for shrub dominated fuel strata. Bootstrap tests of equivalence did not reject the null hypothesis of dissimilarity.

**Figure 2.**
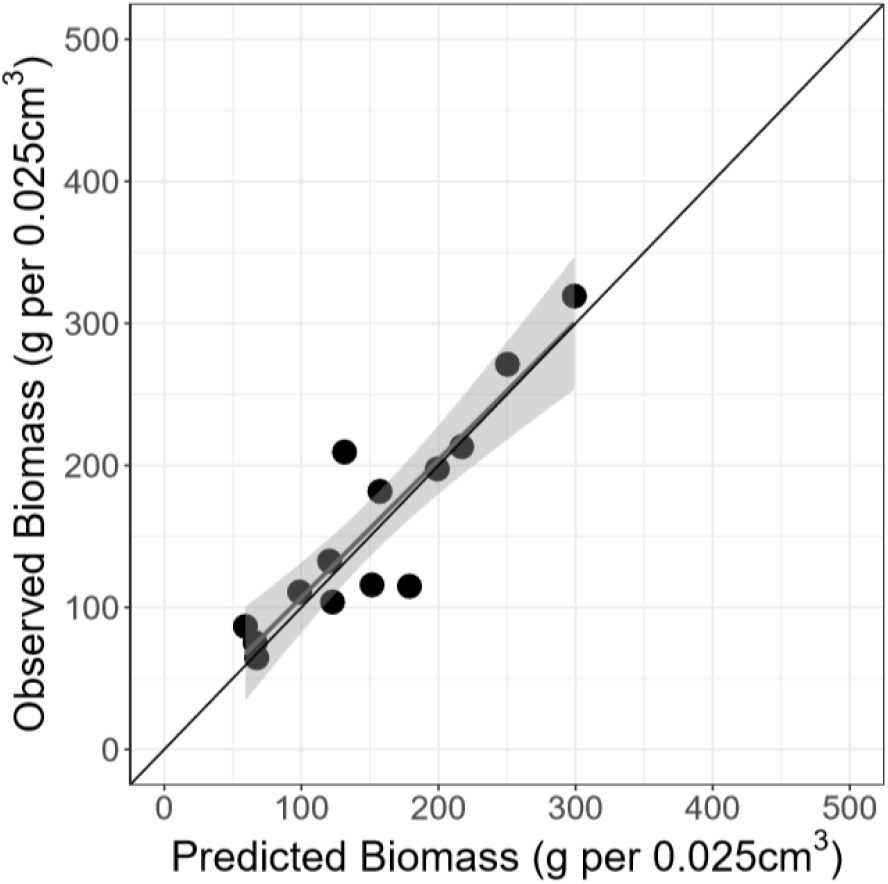
Predicted fuel biomass of the 0-10 cm stratum using the multiple linear model (maximum porosity for 0-10 cm and 10-20 cm) in relationship to the observed fuel biomass for the same stratum.

**Figure 3.**
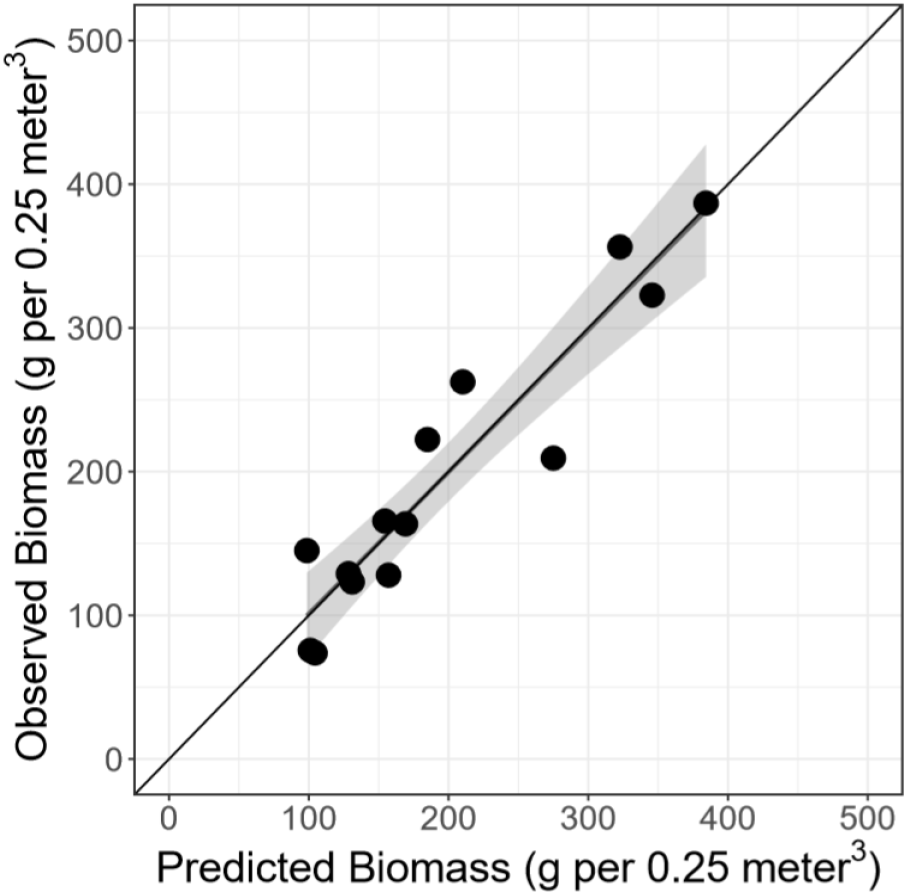
Predicted fuel biomass in relation to observed fuel biomass using the combined estimates of the porosity and surface area models with total observed fuel mass.

### Total Biomass using a Mixed Model

The combined model for total biomass (the sum of the porosity and surface area-based predictions of fuel mass) had a strong relationship with total observed fuel mass including coarse woody debris (Adj. R^2^ = 0.91, **Fig. 4**). This model yielded an error of 16.5% (RMSE=32.64 g) and bootstrap tests of equivalence rejected the null hypothesis of dissimilarity (*P* = 0.025).

**Figure 4.**
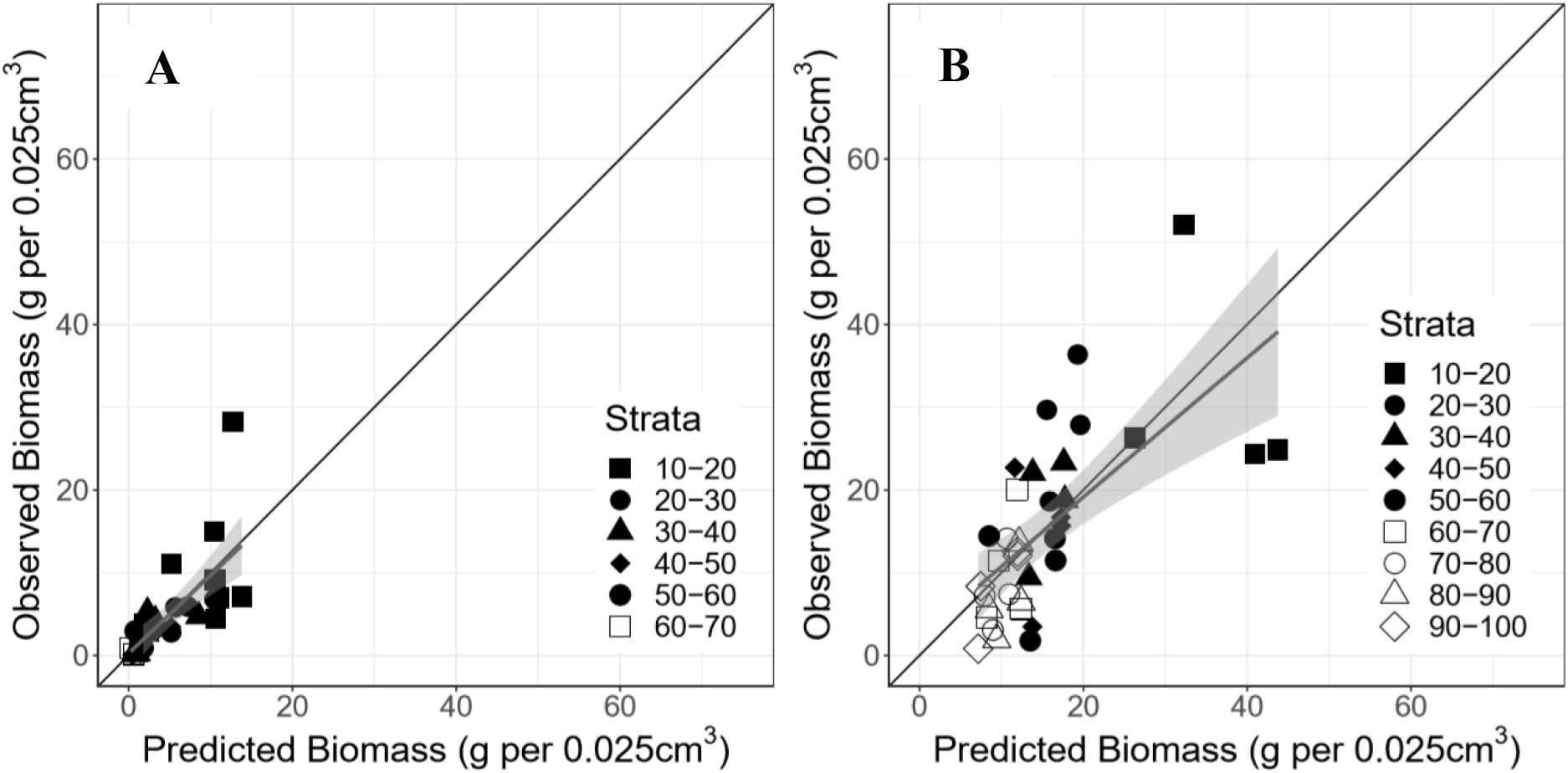
Predicted fuel biomass from surface area of the grass plus forb-dominated plots (A) and shrub-dominated plots (B) with observed fuel mass for all strata.

## Discussion

In this study, we demonstrated the integration of a new approach for measuring three-dimensional wildland fuels in the field with coincident and highly resolved TLS-based fuel parameters. To our knowledge, this study represents the first effort at directly comparing 3-D field data with 3-D TLS data for surface and ground fuels and one of the first for describing fine-scale distributions of biomass three dimensionally in surface fuel layers.

We quantify fuel mass for the 0-10 cm surface fuels using a new fuel metric – porosity. The porosity metric was created because ground layer fuels have been the most difficult to characterize with TLS, yet illustrate the most available fuel mass. For our site, this stratum of fuel accounts for 76.8% (stdev = 21.5%) of the available fuel mass in measured plots and the fuel stratum most likely to contribute to active combustion in prescribed fire environments (Ottmar et al. 2016b). We also quantify fuel features for height strata from 10-100 cm using 10-cm^3^ estimates of surface area that represent a compendium of grasses, forbs, and shrub type fuels. Collectively, these elements affect nearly all aspects of surface fire behavior in the longleaf pine ecosystems sampled in this study, including fluid flow of air that influences fire spread and intensity. Finally, we integrated these independent estimates of fuel mass into a combined model with strong correlations (Table 1).

**Table 1.**
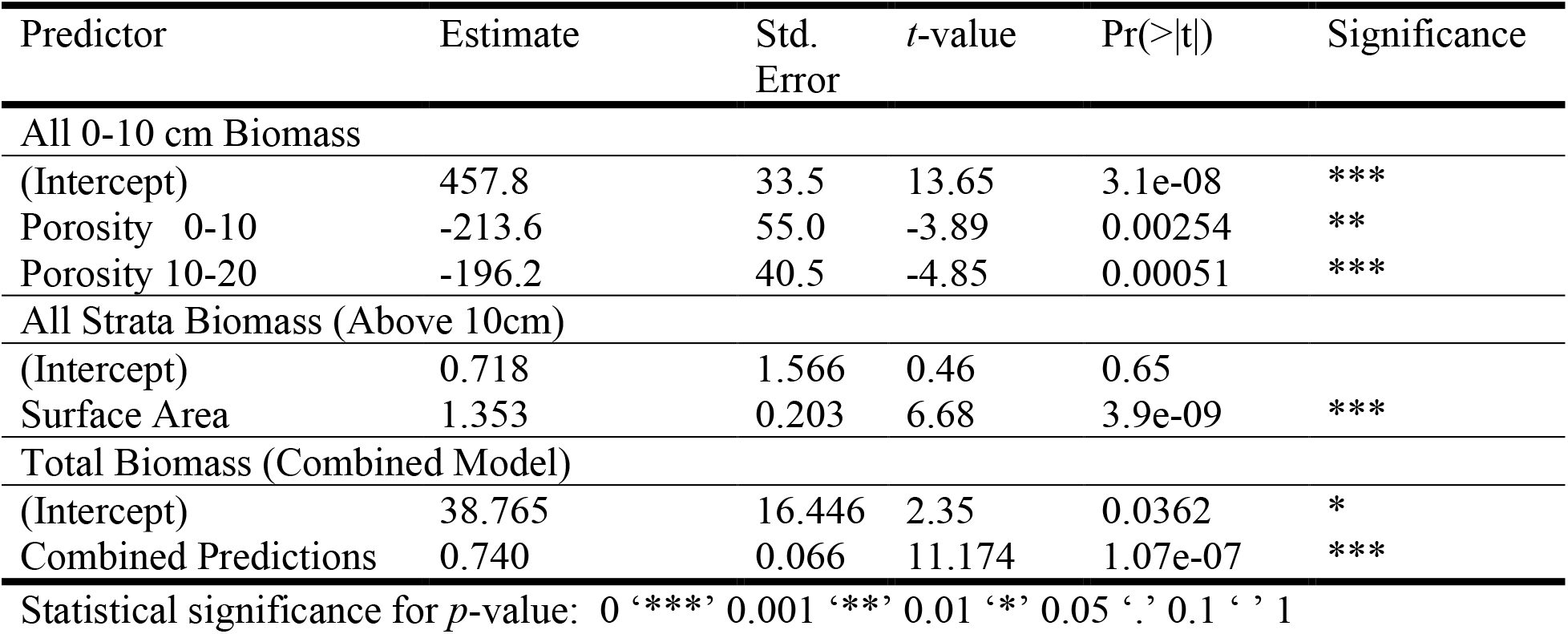
Linear regression model coefficients predicting pre-fire fuel mass from porosity, surface area, and combined models described in this study.

### Estimates of Fuel Mass from Occupied Volume

We expected that occupied volume as an independent metric would explain more variability in the fuelbed stems because of successful execution of a similar model in the RxCADRE experiments at Eglin Air Force Base, FL, USA (Ottmar et al. 2016a). Although Rowell et al. (2016) found strong relationships between voxel-based occupied volume and two-dimensional fuel mass from clip plots within grasslands, this current study shows that the 3D voxel field sampling estimates of occupied volume were not as strongly related to total fuel mass in forested sites. In fact, TLS-based estimates of occupied volume suggest a stronger relationship with measured fuel mass than 2D field estimates. Ultimately, the TLS provided more detailed information on fuel volume distribution than the field estimates. This may be particularly important in the lower strata because of the fine-scale (<100 cm^3^) variability and large amount of mass found in these layers. The main difference between these two studies are the scale at which the field data were collected for the analysis (2D:1m^2^ vs. 3D:0.025 m^3^). Other differences include discrepancies between the Eglin and Pebble Hill sites and their representative species. The primary grass species at Pebble Hill is wiregrass (*Aristida stricta*), as opposed to a broader mix of grasses at Eglin. Shearman et al. (2019) reported that basal volume of the grass tussock was a strong predictor of total aboveground biomass of wiregrass clumps, also describing a local range of biomass at fine scales (both live and dead) at approximately ±200 g. From our results, we hypothesize that biomass density is more variable than suggested by occupied volume estimates. Therefore, there is a need to evaluate these fuels at a finer grain and with a more sensitive metric, such as our TLS-derived porosity, than overall occupied volume. Our results also suggest that the occupied volume estimates from TLS is equally insensitive to variability in the litter layer. As previously mentioned, this lowest 10-20 cm stratum accounts for most of the fuel in these systems and often least represented using remote sensing.

### Estimates of Fuel Mass by Strata

The segmentation of the plots into 10-cm strata served as an optimal approach to assess within plot variability, particularly when the TLS was used to describe porosity in the lower litter layers. Using the 3D field sampling protocol (Hawley et al. 2018), all plots exhibited 100% occupancy for all the 0-10 cm strata and voxels therein. The porosity metric was therefore developed to characterize variation in this compact stratum. The porosity metric could be used to represent a pseudo-packing ratio value (Rothermel 1972), where the range of open space within the 10-cm^3^ voxels increased sensitivity to fuelbed mass variability. The use of both the 0-10 cm and 10-20 cm maximum porosity proved advantageous, because from the field data, we found that litter and downed debris is predominantly found up to 20 cm (unpublished data). Moreover, the use of ‘maximum’ porosity provided a broader range of porosity not encompassed in the mean porosity metric, though the models for both maximum and mean porosity performed equally well, with a slightly steeper regression slope for mean porosity. Moreover, we found that porosity was a strong predictor of litter fuel mass across a range of fuel types including pine needles, deciduous litter, pine cones, and coarse woody debris (unpublished data). Leaf litter in particular is difficult to characterize explicitly. For example, Hudak et al. (2016) calculated fuel mass at 5m × 5m grain size from airborne laser scanning and identified fine fuels as being most problematic to characterize due to occlusion of the forest floor and the coarse sampling grain of airborne laser systems. Newer TLS systems overcome this problem through advanced processing capabilities that provide for more precise--and likely more accurate--estimates to distinguish between true ground and the surface litter layer.

Surface area was more variable as a predictor of fuel mass most likely due to its tendency to depict the outer hulls of the sub-voxel fuel elements and not specifically describing the intra canopy dynamics of stems and twigs that have differing fuel densities than the outer leaves. This effect was most pronounced at the lower strata because TLS-based surface area generally underestimates observed fuel mass, while the inverse occurs at the higher strata (Fig. 3).

## Conclusions

Increasingly, it is recognized that advances in our understanding of how wildland fire drives ecological fire effects is dependent on mechanistic connections of fuels, energy release, and future vegetation (Mitchell et al. 2009, O’Brien et al. 2018). The application of novel 3D sampling techniques to high resolution TLS data in this study represents a major advancement in representing critical variation of fuel characteristics that drive these fire-vegetation feedbacks. The ability to predict both the distribution of mass and 3D structure of surface fuels will allow emerging fire behavior models (Linn et al. *In review*) to create more realistic scenarios for fire planners and fire managers to predict desired fire effects. A next step is to separate these 3D fuel beds into surface or ground fuels (litter, coarse woody debris, grasses) and shrub layers because these two types of fuel affect fire propagation and intensity differentially. This is because fuel elements in the litter stratum are comprised of combustible materials that carry fire and the shrub objects generally represent objects that produce wind flow drag. In CFD fire behavior models such as HIGRAD\FIRETEC, shrubs are modeled as independent objects that are treated as small trees with a homogenized mesh architecture. The advancement in understanding the variability of fuel mass of shrubs at fine-scales (10 cm^3^) allows for improvements in representing understory features from the object perspective, thus adding heterogeneity to the object models used in the CFD fire behavior models. The inclusion of more heterogeneous fuel properties into these CFD models is expected to improve characterization of fluid flow that affect fire intensity and spread, as well as resulting fire effects (Loudermilk et al. 2018; O’Brien et al. 2018).

## Acknowledgements

We acknowledge Tall Timbers Research Station for their support, including Kevin Robertson and Pebble Hill Plantation, and for hosting the Prescribed Fire Consortium, an assembly of researchers and practitioners with a common goal to support prescribed fire applications and advance prescribed fire science. This study was funded in part by the US Department of Defense Strategic Environmental Research and Development Program (projects: RC-2243, 2643), which has sparked ongoing work (projects: RC-1329, 1064, 1119, 2640, 2641). This work was also funded in part by the National Center for Landscape Fire Analysis at the University of Montana. This work was also supported by the US Department of Agriculture (USDA) Forest Service National Fire Plan. We acknowledge the Fire and Environmental Research Applications Team of the USDA Forest Service, Pacific Northwest Research Station, including Roger Ottmar and the cooperative research through the University of Washington, School of Environmental and Forest Sciences, including Susan Prichard. We acknowledge Rodman Linn, Los Alamos National Laboratory, and Nicholas Skowronski, USDA Forest Service, Northern Research Station, for insight and feedback throughout the years. We acknowledge the Southern Research Station, Forestry Sciences Laboratory, Athens, GA for their support.

